# Decisive Role of Polymer-BSA Interactions in Biofilm Substrates on ‘Philicity’ and EPS Composition

**DOI:** 10.1101/2021.02.26.433004

**Authors:** Suparna Dutta Sinha, Madhumita Choudhuri, Tania Basu, Debkishore Gupta, Alokmay Datta

## Abstract

Formation of extracellular polymeric substances (EPS) is a crucial step for bacterial biofilm growth. Dependence of EPS composition on the growth substrate and the conditioning of the latter is thus of primary importance. Here, we present results of studies on the growth of biofilms of two different strains each, of the Gram negative bacteria *Escherichia coli* and *Klebsiella pneumoniae*, on four polymers used commonly in indwelling medical devices – Polyethene, Polypropylene, Polycarbonate, and Polytetrafluoroethylene immersed in Bovine Serum Albumin (BSA) for 24 hrs. The polymer substrates are studied before and after immersing in BSA for 9 hrs and 24 hrs, using contact angle measurement (CAM) and Field Emission Scanning Electron Microscopy (FE-SEM) to extract, respectively, the ‘philicity’ (defined as *ϕ* ≡ *sin* (*θ*-90°), where *θ* is contact angle of the liquid on the solid at a particular temperature and ambient pressure) and spatial Hirsch parameter *H* (defined from the relation, *F(r)* ~ *r^2H^*, where *F(r)* is the mean squared density fluctuation at the sample surface). *H* =, <0.5 or >0.5 signifies no correlation, anti-correlation, and correlation, respectively. The substrates are seen to transform from large hydrophobicity to near amphiphilicity with the formation of BSA conditioning surface layer, and the *H*-values distinguish the length scales of ~ 100 nm, 500 nm, and 2000 nm, with the anti-correlation increasing with length scale. Biofilms grown on the BSA-covered surfaces are studied with CAM, FE-SEM, Fourier Transform Infrared (FTIR) and Surface Enhanced Raman Spectroscopy (SERS). Most notably, the *ϕ*-values are *independent* of the bacterial species and strain but *dependent* on the polymer, as is also shown strikingly by both types of spectra, while *H*-values show some bacterial variation. Thus, the EPS composition and consequently the wetting properties of the corresponding bacterial biofilms seems to be decided by the interaction of the conditioning BSA layer with a specific polymer used as the growth substrate.

## 1. Introduction

Biofìlms form the collective mode of existence of bacteria apart from the isolated or planktonic mode. Though they were detected by Leeuwenhoek in the earliest instance of microscopy, basic understanding about biofilms, especially their striking levels of organization and properties, has emerged only recently [1–3]. On one hand, they have proved to be highly beneficial in providing sustenance in both land and aquatic environments by maintaining the bacterial population networks at soil and water surfaces [4–7]. On the other hand, they pose grave medical threats, by forming pathogen organizations within the physiological environment on indwelling medical devices and biomaterial implants [8–11], which form a significant part of healthcare associated infection worldwide [12–14]. Biofilms in clinical samples, which have been much less presented outside dental plaques, are always found to develop attached to a surface unless dislodged from the site of infection [15]. Hence, the specifics of the surface structure and compositions are some of the key determinants of biofilm growth and development as they decide the bacteria-surface forces.

Bacteria organize themselves into biofilms through the formation of extra-cellular polymeric substance or EPS [16]. This is highly heterogeneous and poorly defined [17], with polysaccharides as a primary/base constituent accompanied with a variety of proteins, lipids [18,19], and smaller organic molecules serving different functions in the biofilm. The composition of EPS is known to vary among biofilms of different bacterial species, and maybe even among biofilms of the same species under different growth conditions [20,21]. The composition of EPS is known to decide the functionality of the biofilm; hence the control of this composition can effectively control its pathogenic activity [22,23]. However, what is most important is that in the absence of the EPS or in the event of its composition being suitably changed, biofilm growth itself may be stopped. The indepth knowledge about the contribution of albumin in surface modification of biomaterials and the manner in which the biofilm bacteria respond to it through substrate-specific variation of the chemical constituents of the EPS matrices will not only enhance the knowledge of the initial stages of biofilm formation on medical devices, but will also be crucial in developing non-invasive strategies of biofilm control.

Indwelling medical devices are in general composed of a few common polymers such as Polyethene, especially high density polyethene or HDPE (in knee and hip implants), Polypropylene or PP, Polycarbonate or PC, and Polytetrafluoroethylene or PTFE. Major reasons for using these polymers are their hydrophobicity, high degree of chemical inertness and mechanical strength and lightness (especially in case of HDPE). However, the devices in contact with physiological fluids, tend to get adsorbed with the proteins from plasma and other body fluids within nanoseconds of its introduction [24,25], much before the arrival of the microbes or tissue cells. Hence the pathogens and the surrounding tissue cells, which appear at the site of introduction of the medical device-much later, view the ‘surface with the adsorbed protein layer’ as the ‘original foreign surface’ instead of it being only a ‘conditioning layer’ [26]. This conditioning layer, especially albumin has the propensity to alter the hydrophobicity and chemical sterility [27] of the surface of the medical devices in a manner that may finally lead to septicaemia or irreversible device-associated infections through the formation of biofilms. Since biofilms exhibit very low susceptibility to antibiotics, it is imperative to understand the manner in which albumin modulates the polymer surfaces to enhance of biofilm growth.

In this communication, we present the results of our investigations regarding two issues, in two separate sections - Part I: how the interaction between serum albumin and the polymers mentioned above may change the surface property, specifically the wetting property of the polymer biomaterials, and Part II: how the interactions between these modified surfaces and the major organic components of biofilms grown on them change the wetting property and, more important, the composition of the EPS. Instead of HSA we have used the more standardised and reproducible bovine serum albumin (BSA) with essentially the same composition. Clinical and American Type Culture Collection or ATCC strains of Gram-negative bacteria *Escherichia coli* or *E. coli* (EC and EA respectively) and *Klebsiella pneumoniae* or *K. pneumoniae* (KC and KA respectively) were used as the biofilm forming bacteria. We have found that (1) while BSA-polymer interactions vary from polymer to polymer, the trend is to reduce the hydrophobicity to a value where the surface can be wetted by water (or other hydrophilic substances) as well as by hydrocarbons (or other hydrophobic substances), i.e. modulation of the hydrophobic substrates to amphiphilic substrates and (2) these amphiphilic substrates give rise to EPS that vary in composition from polymer to polymer but very similar to each other for the different bacterial strains used.

## 2. Part I: Bovine Serum Albumin-coated Polymer Substrates

### 2.1 Materials and Methods

#### 2.1.1 Adsorption of BSA on Polymer samples

PP is used in a variety of catheters except urinary catheters, while HDPE is widely used in orthopedic implants. The PTFE balloon catheter is especially designed for short- and medium-term urinary drainage, while other catheters are often lined with PTFE to reduce friction and to ensure that other devices can pass through with ease. PC is widely used in indwelling medical devices, where there is a requirement of toughness, rigidity and visual clarity, including minimally invasive surgical devices. Hence the above widely used polymers have been used in our experiments as substrates for growing biofilms.

Commercially available clinical grade (biocompatible) PP, HDPE, PTFE and PC chips having machined finish were obtained in square configuration (10 mm X 10 mm) from Plastic Abhiyanta Ltd, India. The water used in all stages of our experiments was of HPLC grade (Lichrosolv) from Merck, India. Tris buffer was obtained from Sigma-Aldrich, USA while BSA was from MP Biomedical Ltd, USA. The polymers chips were initially cleaned in an ultrasonic cleaner, rinsed with water, blow dried, and preserved in a vacuum desiccator, ready for carrying out adsorption [28]. BSA was mixed in the proportion 0.5 mg/ml with buffer solutions of pH 7.4 and left for a week with intermittent mixing to dissolve the BSA completely. BSA solutions were taken in four sets of separate glass vials, each containing a single chip of each category (PP, PC, PTFE and HDPE) for 9 hrs. The similar procedure of adsorption was repeated for a fresh set of polymer chips, for 24 hours. This stage was conducted in triplicate for both cases. The BSA was left to adsorb on the polymer samples for a certain period of time (9 hours and 24 hours), to simulate a static situation, where a biomaterial placed in a physiological environment gets adsorbed with plasma proteins. This time is hence referred to as the ‘adsorption time’. After the stipulated period, the chips were removed from the protein solutions, rinsed with water, and finally blow-dried and preserved in a desiccator ready for growing biofilms. The chips obtained, possessed different amounts of BSA adsorbed on them, termed as the ‘conditioning layer’. They could not be sterilized further, to prevent the denaturation of the adsorbed protein and were preserved for growing biofilms on them.

#### 2.1.2. Contact Angle Measurement (CAM)

The contact angles of water on the different polymers with and without the ‘conditioning layer’ of adsorbed BSA (for 9 hours and 24 hours) were measured using, static sessile drop method [29]. The angle formed between the solid and the liquid interface was measured using a microscope optical system for capturing and Image J was used to determine the contact angle. All contact angle observations were performed in triplicate. [See SI]

#### 2.1.3. Field Emission-Scanning Electron Microscopy (FE-SEM)

The dried polymer chips with and without different proportions of ‘conditioning layer’ were sputter-coated with a 3-nm-thick conductive layer of gold. FE-SEM measurements were conducted on them, at 2.0–10 kV with a Inspect F50 Microscope (FEI Europe BV, and The Netherlands; FP 2031/12, SE Detector R580). We have used Image J again for analysis of the FE-SEM images.

#### 2.1.4. Philicity and Hirsch Parameter

##### Philicity

We introduce a term ‘philicity’ to quantify the wetting characteristics of a surface by a liquid. We define it as *ϕ* ≡ *sin* (*θ* - 90°), where *θ* is contact angle of the liquid on the solid at a particular temperature and ambient pressure. For water, *ϕ* <0 implies the substrate to be hydrophilic, *ϕ* > 0 means it is hydrophobic, while *ϕ* = 0 stands for a substrate that is neither and maybe termed ‘amphiphilic’.

##### Hirsch Parameter

The FE-SEM images of the samples yield the mean square intensity fluctuations at different length scales. This spatial variation of intensity along a typical line in each of the observed images can be assumed to follow the spatial variation of sample density under observation. The mean square (ms) intensity fluctuation [30], *F*(*r*), *r* being the separation between points along typical lines, is given by

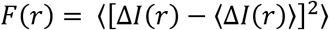

where *I* is image intensity at point (*x, y*), Δ*I*(*r*) = *I*(*x,y*) — *I*(*x*_0_,*y*_0_), where (*x*_0_, *y*_0_) is the reference point and 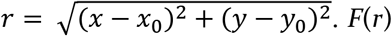 thus represents the mean squared density fluctuation at the sample surface.

The spatial Hirsch parameter *H* is defined from the relation, *F*(*r*) ~ *r*^2H^. This Hirsch parameter divides any spatial pattern into three groups, correlated, uncorrelated, and anticorrelated, accordingly as its value is more than, equal to, or less than 0.5, respectively. Again, this signifies the corresponding presence of attractive, zero, or repulsive forces in the system, with the deviation of the value of *H* from 0.5 serving as a measure of the strength of the force. This has proved to be a very useful technique to gauge forces is soft and complex matter where a direct measurement is extremely difficult [31–33].

### 2.2. Results

#### Scanning Electron Microscopy and Hirsch Parameter

Figure 1 represents the FE-SEM images of some of the bare polymer surfaces (Figures 1(a) and 1(b)) and the surfaces of the same polymers immersed for 24 hrs in BSA(Figures 1(c) and 1(d)). The mean square density fluctuation (MSDF) *F*(*r*), and the *H*-values from the log-log plots of *F*(*r*) versus *r*, extracted from these images are also presented in this figure (Figures 1(e)-1(h)). It is clear that the *H*-values have different constant values over three different length scales, around 100 nm, 500 nm, and 2000 nm or 2 μm. It is interesting to note that the first length scale matches with the contour length of BSA, which will be attained by a BSA molecule fully stretched on the substrate surface, while the last matches with the dimensions of most bacteria.

**Figure 1:**
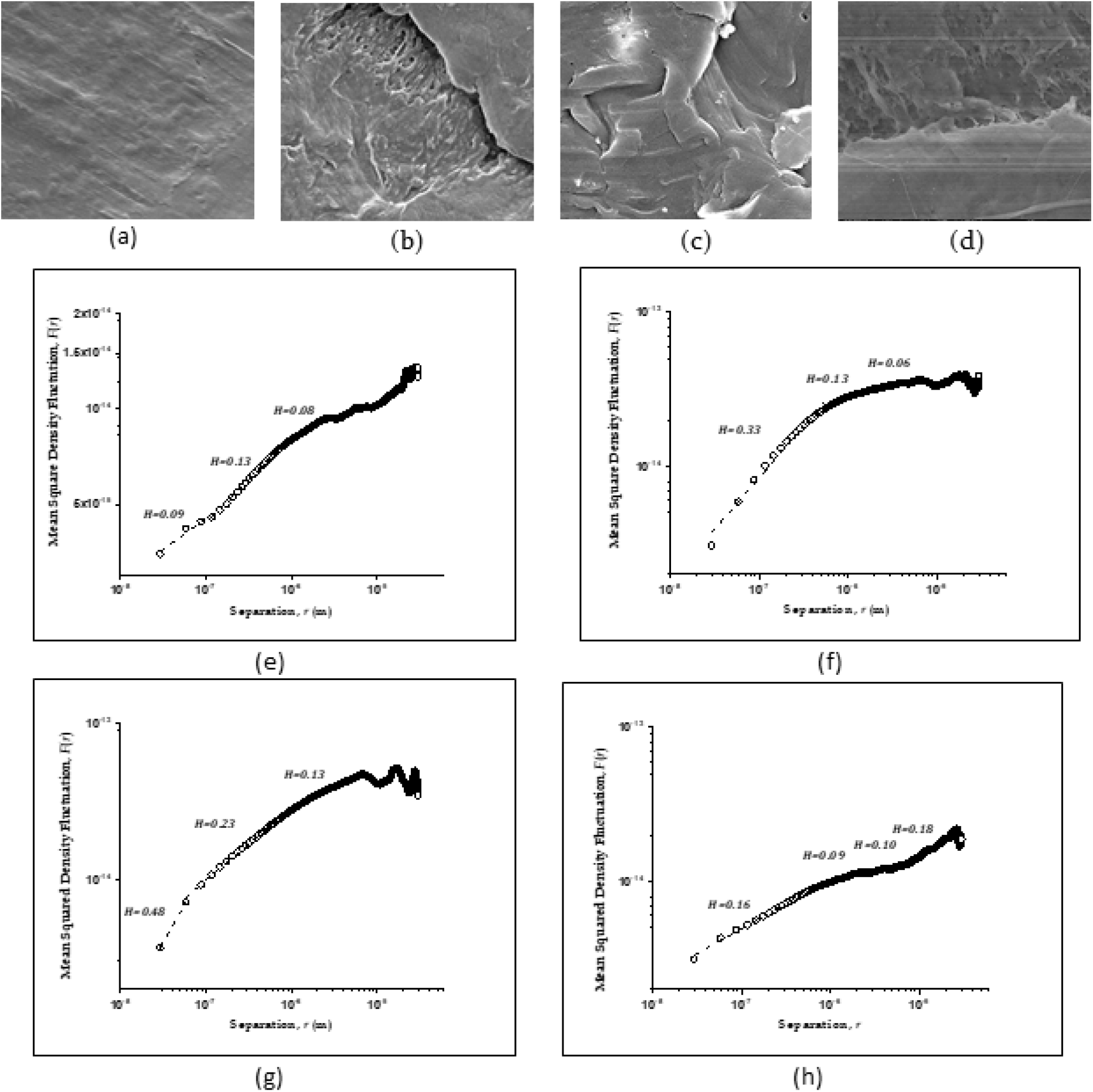
Representative Scanning Electron Microscopic (FE-SEM) images and mean squared density fluctuation (MSDF) of bare and Bovine Serum Albumin (BSA) coated polymers. (a)-(d): FE-SEM images, (e)-(h): MSDF, *F*(*r*) versus *r* (in m) extracted from (a)-(d). Bare (a) High Density Polyethene (HDPE), (b) Polytetrafluoroethylene (PTFE) and 24 hrs BSA immersed (c) HDPE and (d) PTFE. (e) MSDF from (a), (f) MSDF from (b), (g) MSDF from (c), (h) MSDF from (d). Log-log plot of data are shown in open circles. Fits with *F*(*r*) ~ *r^2H^* are shown in dashed lines, with values of the Hirsch parameter *H* shown beside fitted segments.

#### Philicity and Hirsch Parameter

Figure 2 summarizes the results obtained from CAM measurements (SI) on bare HDPE, PTFE, PP, and PC, and on these polymers kept immersed in BSA for 9 hrs and for 24 hrs and compares these with those obtained from FE-SEM measurements of these polymers after 24 hrs of BSA immersion. It shows the values of ‘philicity’ from the CAM measurements and *H*-values from the FE-SEM measurements. The major common trend from ‘philicity’ values is a change from hydrophobicity for bare and for 9 hrs immersed polymers to near amphiphilicity, 0.05 > *ϕ* > −0.05, for 24 hrs immersed polymers. For *H*-values, the trend is anti-correlation at all length scales and for all BSA immersed polymers, other than for HDPE at 100 nm length scale where there is no correlation (*H* ~ 0.5). It is also seen that the anti-correlation increases with the length scale. This indicates the presence of short and long-range repulsive forces in the BSA covering the polymer surfaces, except for HDPE, where at the short-range the repulsive and attractive forces balance each other.

**Figure 2:**
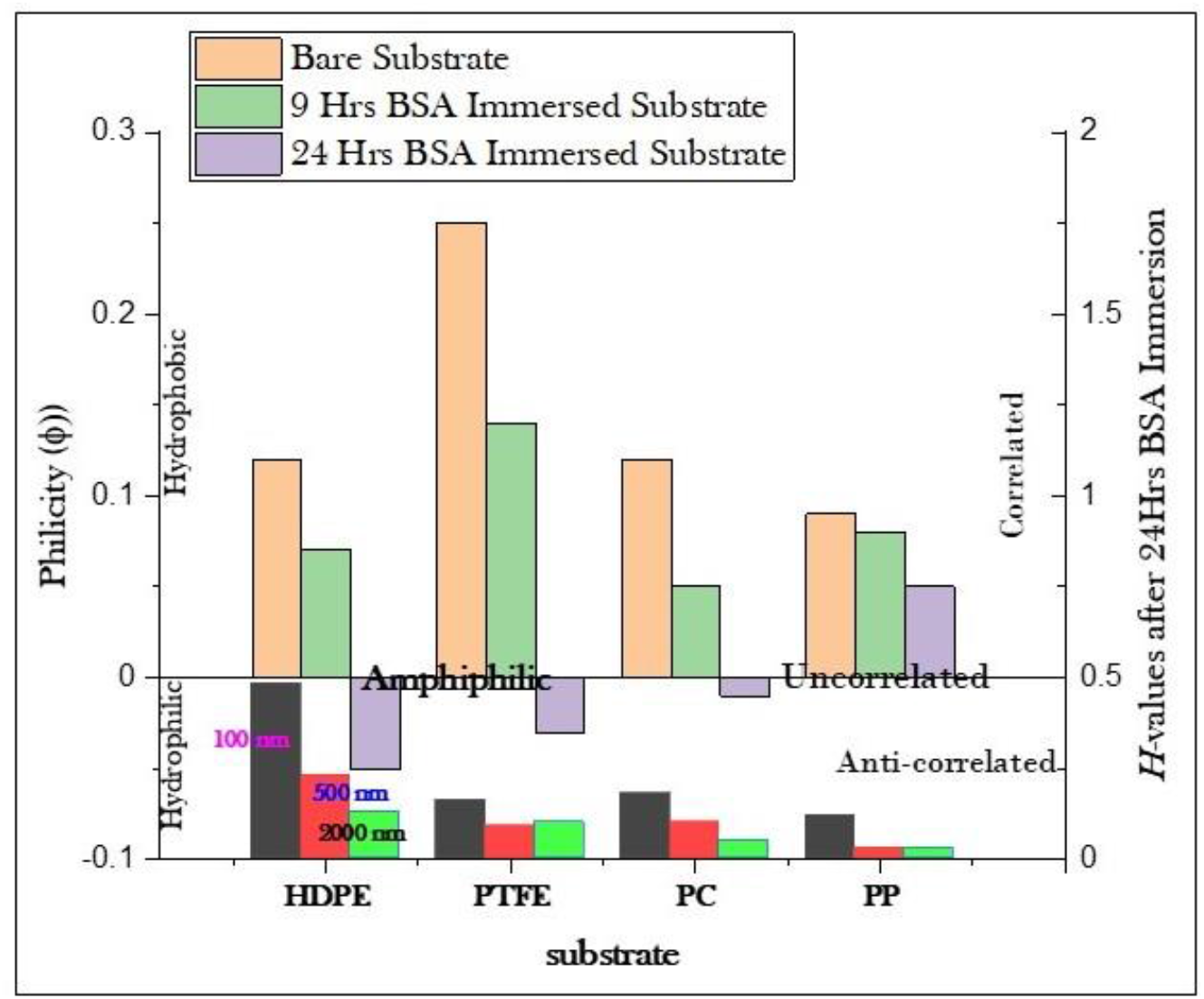
Philicity of the bare polymer substrates, substrates after 9 hrs and after 24 hrs of immersion in BSA, and Hirsch parameter values at different length scales for substrates after 24 hrs of immersion in BSA. Please see text for explanation of terms.

### 2.3 Discussion

A qualitative explanation of these results requires a look at some of the structural aspects of the polymers and BSA. The molecular structures are shown in Figure 3. The presence or absence of polar groups in the polymers and BSA is, of course, related to its hydrophilicity or hydrophobicity, respectively. In particular, a molecule with hydrocarbon or fluorocarbon chains and without any polar group will be strongly hydrophobic, whereas a molecule with a number of polar groups and much more so with dissociable groups will be strongly hydrophilic. When the proportions of the hydrocarbon part and the polar/dissociable part become equal or comparable, the molecule becomes amphiphilic. If a surface is dominated by hydrophobic or hydrophilic molecules it acquires the respective character. If there are either amphiphilic molecules or equal or comparable proportions of hydrophobic and hydrophilic molecules at the surface, the surface becomes amphiphilic.

**Figure 3:**
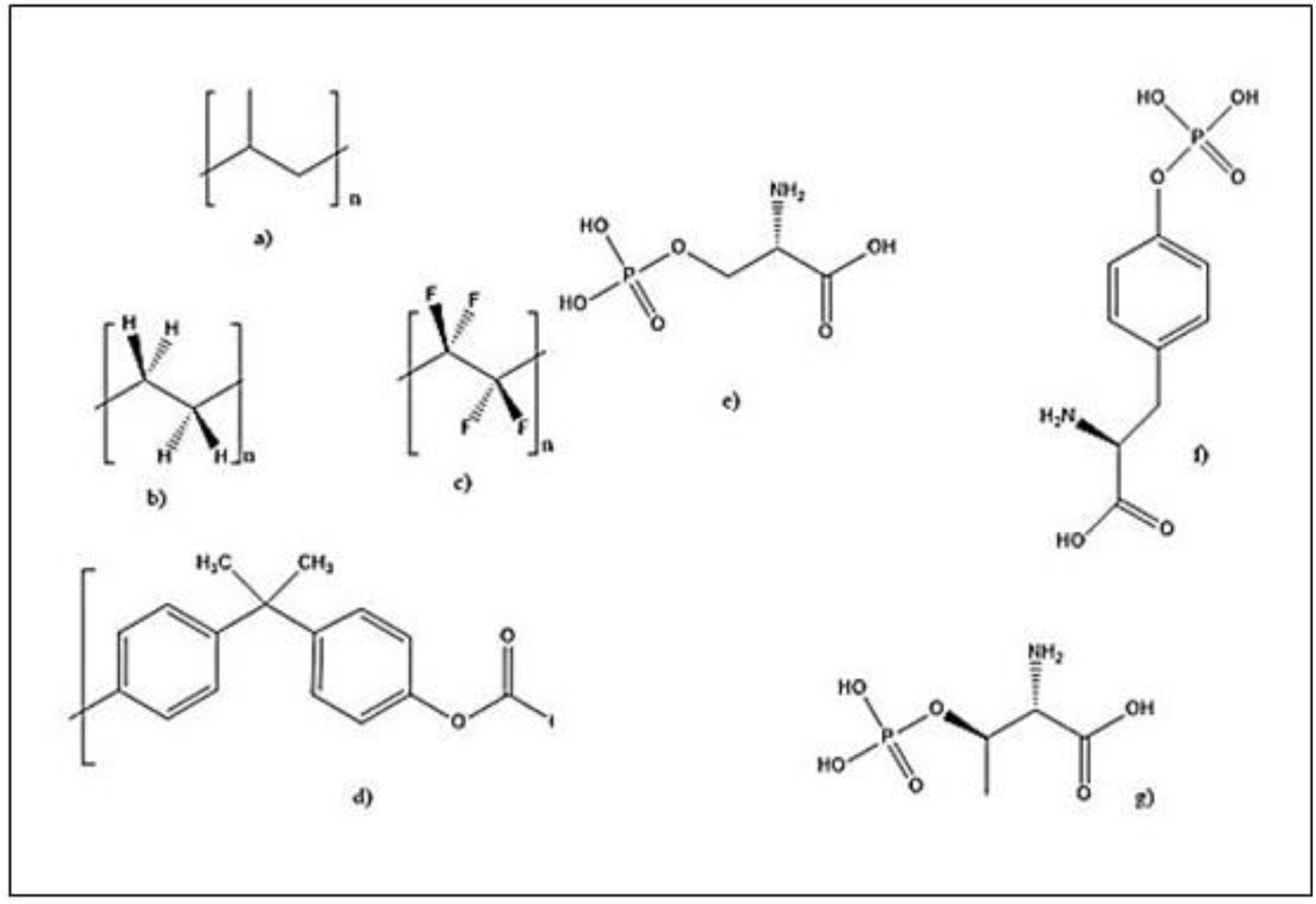
Structures of (a) Polyethene, (b) Polytetrafluoroethylene, (c) Polypropylene, (d)Polycarbonate; Structures of (e) Phosphoserine, (f) Phosphotyrosine, and (g) Phosphothreonine.

Figures 3(a)-(d) show that the polymers have hydrocarbon/fluorocarbon moieties only or dominantly. On the other hand, BSA has a hydrocarbon backbone with disulphides, and a large number of amino acids (Figures 3(e)-(g)) polar and dissociable groups such as carboxyl (CO_2_^-^H^+^), phosphatidyl (PO_3_^2-^H_2_^2+^), and amine (NH_2_) attached. The former explains the hydrophobicity of the bare polymers but the acquired near-amphiphilicity of the 24 hrs BSA immersed substrates calls for more explanation. The most plausible would be the presence of a short-range lipophilic interaction [20] between the hydrocarbon backbone of the BSA molecule and the polymer, which leaves the molecule stretched on the surface with polar/dissociable groups free. This later renders hydrophilicity to the already hydrophobic substrate surfaces and the balance makes the surfaces nearly amphiphilic.

The trend in intermolecular forces is consistent with this explanation since the repulsive force becomes stronger at longer ranges. At shorter ranges the lipophilic attraction between the hydrocarbon backbone of BSA and the polymer competes with the repulsion between the dipolar groups of BSA fixed on the substrate surfaces. At longer ranges, only the dipole-dipole repulsion remains [32].

## 3. Part II: Bacterial Biofilms on BSA-coated Polymer Substrates

### 3.1 Materials and Methods

#### 3.1.1 Bacterial Species

The clinical strains of Gram-negative bacteria, *E. coli* and *K. pneumoniae*, were obtained isolated from fatally ill patients, through the central catheter system, of The Calcutta Medical Research Institute, Kolkata. The Gram-negative strains of E coli ATCC 25922 and Klebsiella pneumoniae ATCC 700603 were from American Type Culture Collection.

#### 3.1.2. Cell culture and preparation

The bacteria were cultivated at 150 rpm in standard Luria-Bertani (LB) medium (10 g of tryptone, 5 g of yeast extract, and 10 g of NaCl per 1 L of deionized water, pH adjusted to 7.2 and sterilized at 121 °C for 20 min). The cultivation temperature for *E. coli* and *K. pneumoniae* was 37 °C. Cells were harvested in the stationary phase after 24 hrs cultivation. The bacteria cells were collected by centrifugation (3,000 rpm, 4°C, 10 min) and washed three times in 165 mM phosphate buffer saline (PBS, composition 1.093 g Na_2_HPO_4_, 0.276 g NaH_2_PO_4_, and 8.475 g NaCl in 1 L deionized water, pH adjusted to 7.2 and sterilized at 121°C for 20 min) to remove the residual LB medium. Bacterial cells were resuspended in PBS to a concentration equivalent to an optical density at 600 nm (OD 600 nm) of about 0.2. The suspension was then used for cell adhesion and biofilm cultivation immediately.

#### 3.1.3. Growth of biofilms

The protocol for biofilm growth [28] was followed with the four sets of polymer chips with BSA adsorbed on them for 24 hours, and three samples of each type were placed in separate wells of a 24 well tissue culture plate with identical growth conditions. Biofilm were simultaneously grown on three samples of each type of bare polymer (without BSA) - following the same protocol. After the completion of the 7-day growth period, the polymer chips were aseptically removed and washed thrice with phosphate buffered saline (PBS pH 7.2) to remove planktonic bacteria. The chips were then air dried and prepared for Field Emission-Scanning Electron Microscopy (FE-SEM) measurements.

#### 3.1.4 Field Emission-Scanning Electron Microscopy (FE-SEM)

The polymer chips with attached bacterial cells (with or without a conditioning layer of BSA), were covered with 2.5% glutaraldehyde and kept for 3 hrs in 4°C after which they were washed thrice with the phosphate buffer solution. They were then passed once through the graded series of alcohol (25, 50 and 75%, twice through 100% ethanol) each for 10 min, finally transferred to the critical point drier and kept overnight to make them ready for biofilm analysis. The dried polymer chips with biofilms were sputter-coated with a 3-nm-thick conductive layer of gold. To compare the 7-day old biofilms produced by the clinically isolated and ATCC strains of *E. coli* and *K. pneumoniae*, on different substrates, FE-SEM measurements were conducted at 2.0–10 kV with a Inspect F50 Microscope (FEI Europe BV, and The Netherlands; FP 2031/12, SE Detector R580). We have used Image J again for analysis of the FE-SEM images.

#### 3.1.5. Fourier Transform Infrared (FTIR) Spectroscopy

All FTIR spectra were obtained using a Fourier Transform Infrared Spectrometer from Shidzu, having temperature-stabilized DTGS (Deuterated Triglycine Sulfate) detectors optimized for MIR with an enhanced SNR of 15,000:1 peak-to-peak for a 5 second scan. The system had KBr beam splitter for MIR for a spectral range 8,300 – 350 cm^-1^ at a best resolution of 0.4 cm^-1^. The samples were tightly fastened to the sample holder before the measurements and it was confirmed that there was absolutely no space between the two. The spectra of polymer samples with and without BSA, with biofilms attached on them were obtained after running the background scan. The final spectrum in each case was obtained after the background correction.

#### 3.1.6. Surface Enhanced Raman Spectroscopy (SERS)

AgNPs were prepared [34] by reduction of silver nitrate with hydroxylamine hydrochloride at alkaline pH and room temperature [35]. Briefly, 10 mL of silver nitrate (10^-2^ M) was rapidly added, while stirring, into 90 mL of premixed solution containing hydroxylamine hydrochloride (1.67 × 10^-3^ M) and sodium hydroxide (3.33 × 10^-3^ M). Then, the solution was being stirred for 1 min to enable the overall reaction to complete. The AgNP solution was stored in darkness at 4°C. The SERS experiments were done with freshly prepared AgNP solution.

All SERS data were obtained using Lab Ram HR 800 (Horiba Jobin Yvon) spectrometer. The instrument acquired data over a range of 100 cm^-1^ to 3000 cm^-1^ with a 5s exposure time. The laser power was 17 mW, and the operating wavelength of the laser was 632.8 nm. Spectral detection was done using a CCD detector. An Olympus optical microscope was attached to the system and observations were repeated with x10 and x100 objectives, to check for accuracy. All measurements were carried out under identical conditions to enable comparative analysis. 200 μL of prepared AgNP solution was added on the polymer chips with biofilms attached on them, prior to the measurements [36,37]. The samples were then stored in darkness and dried at room temperature. The laser beam was focused on the bacterial layer by applying the Leica × 100 (NA 0.85) objective to a spot of approximately 1 μm diameter.

### 3.2 Results

#### Scanning Electron Microscopy and Hirsch Parameter

Figures 4(a)-(d) present the FE-SEM images of some representative bacterial biofilms grown on substrates that had been immersed in BSA for 24 hrs. While Figures 4(a) and 4(b) show the images of American Type Culture Collection or ATCC strain of *K. pneumoniae* grown on BSA-coated HDPE and PTFE, respectively, Figures 4(c) and 4(d) show images of ATCC strain of *E. coli* grown on corresponding substrates. Figures 4(e)-(h) show the MSDF (log-log plots of *F*(*r*) versus *r*) extracted from Figures 4(a)-(d), respectively, with the *H*-values for the 100 nm, 500 nm, and 2 *μ*m length scales extracted from these plots. From the FE-SEM images, the high coverage of the surfaces by the biofilms is apparent, pointing to the good growth of the latter.

**Figure 4:**
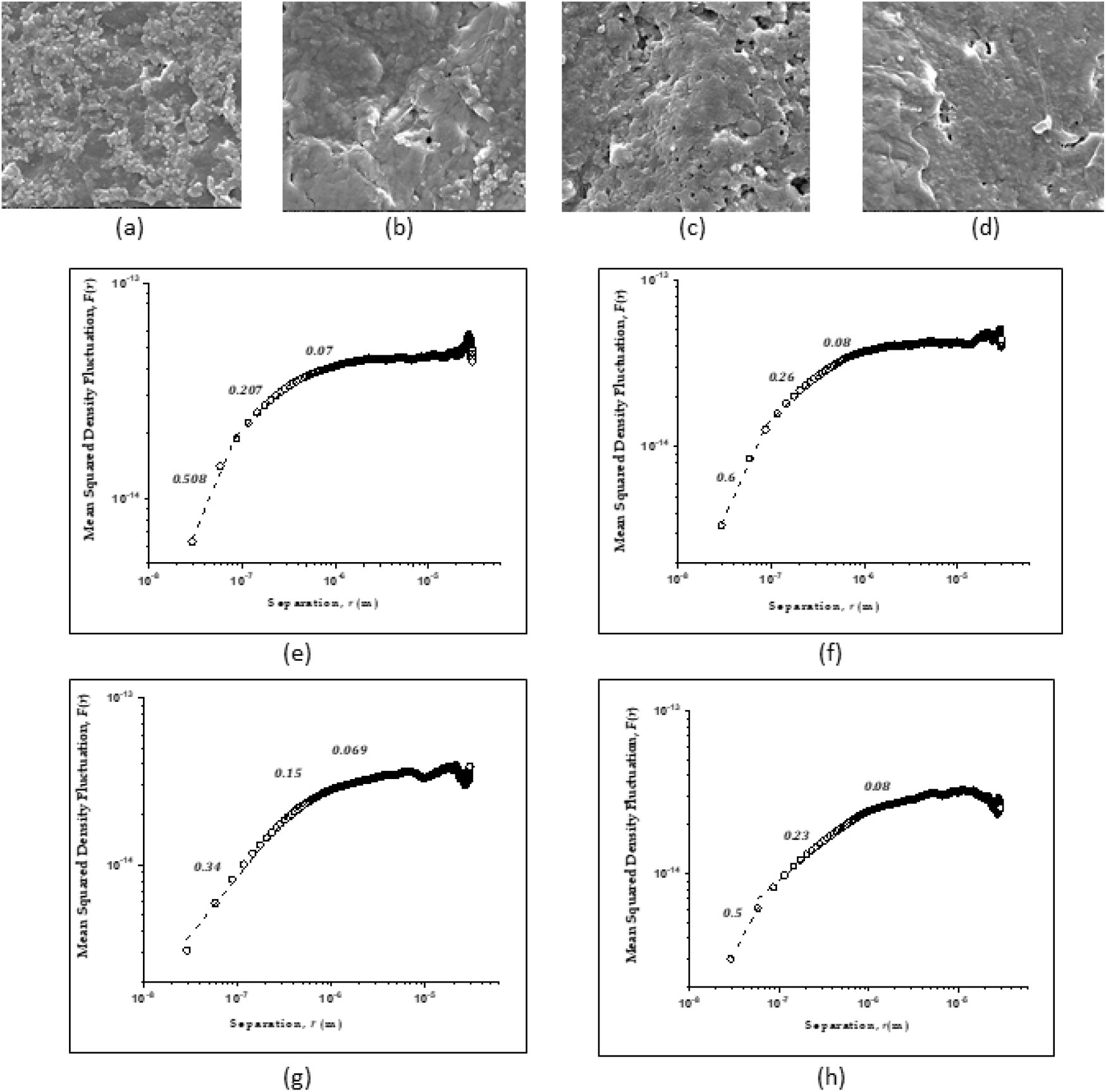
Representative FE-SEM images and MSDF of biofilms grown on BSA (24 hrs) coated polymers. (a)-(d): FE-SEM images, (e)-(h): MSDF, *F*(*r*) versus *r* (in m) extracted from (a)-(d). *Klebsiella pneumoniae* (American Type Culture Collection or ATCC) on BSA-coated (a)HDPE, (b) PTFE and *Escherichia coli* (ATCC) on BSA-coated (c) HDPE, (d) PTFE. (e) MSDF from (a), (f) MSDF from (b), (g) MSDF from (c), (h) MSDF from (d). Log-log plot of data are shown in open circles. Fits with *F*(*r*) ~ *r^2H^* are shown in dashed lines, with values of the Hirsch parameter *H* shown beside fitted segments.

#### Philicity and Hirsch Parameter

Figure 5 summarizes the results obtained from CAM and FE-SEM measurements on biofilms of ATCC and Clinical strains of *K. pneumoniae* (top) and *E. coli* (bottom) grown on the polymers after 24 hr immersion in BSA. From philicity values, the most interesting feature is that these values are independent of the bacterial species and strain but totally dependent on the polymer as the substrate, or in other words, the BSA-polymer interaction has already decided the wetting properties of the biofilms. The next most important feature is the near absence of any change in the ‘philicities’ of the biofilms from those of the BSA-coated polymers shown in Figure 2, which is again consistent with the above proposition.

**Figure 5:**
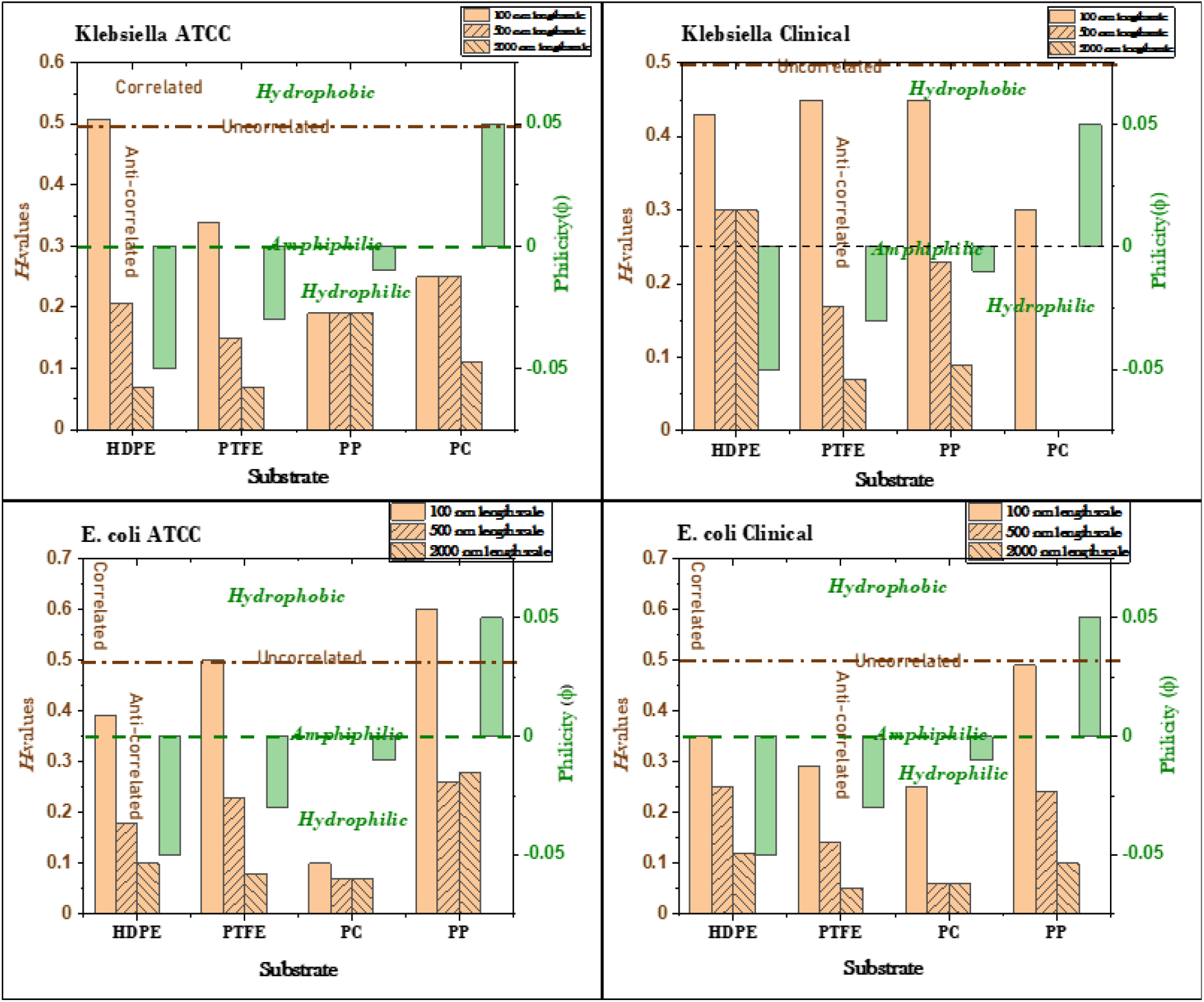
Philicity and Hirsch parameter values at different length scales of the polymer substrates coated with BSA (24 hrs of immersion) and bacterial biofilms on BSA. Top: *Klebsiella pneumoniae* ATCC and Clinical bacterial biofilms; Bottom: *Escherichia coli* ATCC and Clinical bacterial biofilms.

The intermolecular forces on the other hand, as suggested by the *H*-values in Figure 5, show some variation from bacteria to bacteria and also between different strains of the same bacteria. However, the same general trend of short-range attraction competing with repulsion and the stronger repulsion at long range are very much present, consistent with lipophilic and dipolar forces.

#### FTIR and SERS

These expectations are completely borne out by the results of FTIR spectroscopy and of SERS on the biofilms shown in Figure 6 and Figure 7, respectively. The tentative assignment of some of the major peaks in these figures and the probable source of these assigned groups are given, respectively, in Table 1 and Table 2. Both types of spectra again show, strikingly, that the spectra are almost identical for the different species and strains of bacteria on the same polymeric substrate, while they are completely different for the same species and strain of bacteria on different polymers. This establishes the fact that the composition of EPS generated by the biofilm on a specific BSA-coated polymeric substrate is decided by the specific BSA-polymer interaction.

**Figure 6:**
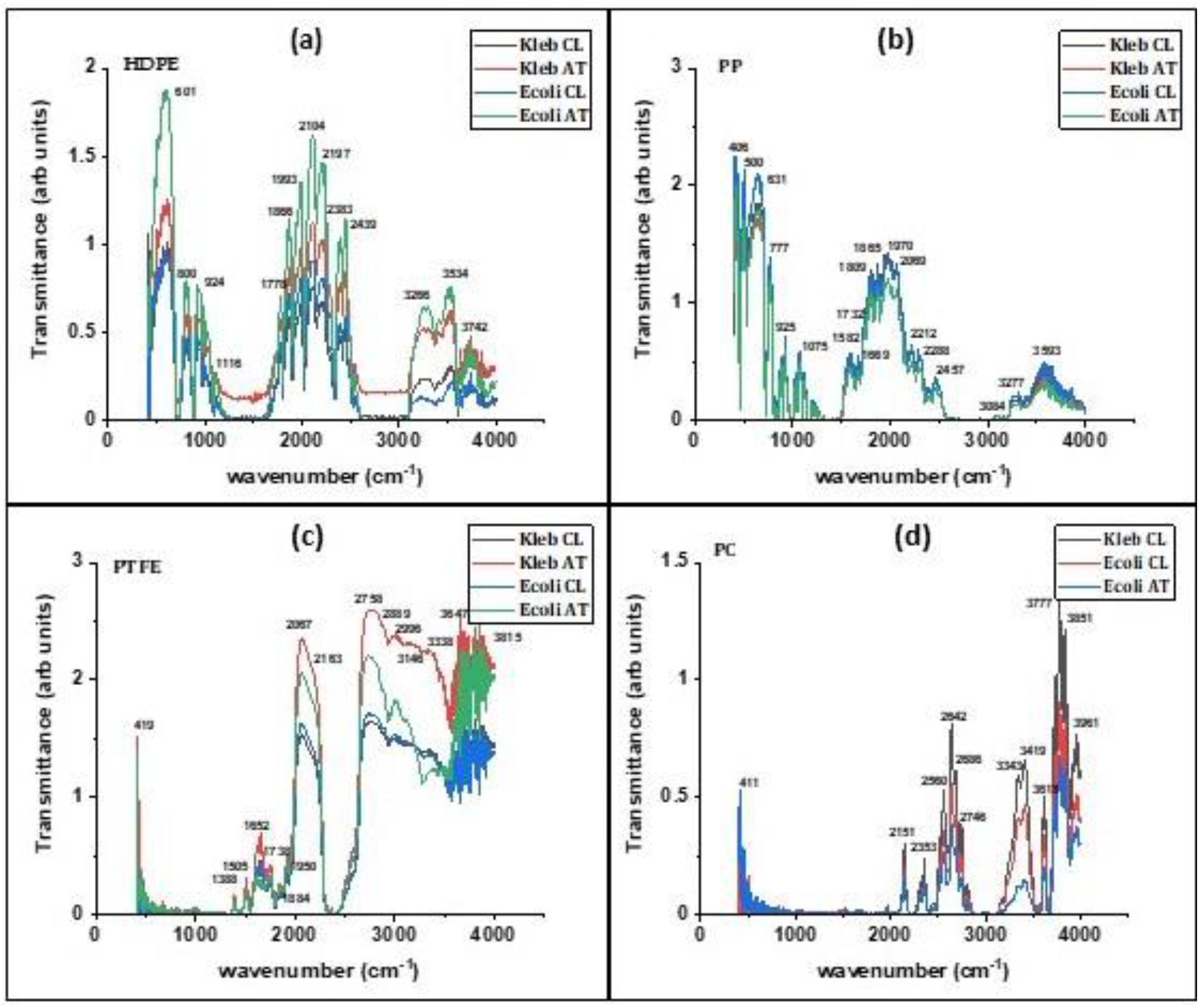
Fourier Transform Infrared Spectra of the bacterial biofilms. *Klebsiella pneumoniae* ATCC and Clinical, and *Escherichia coli* ATCC and Clinical bacterial biofilms grown on (a) HDPE, (b) PP, (c) PTFE, and (d) PC. Prominent peak positions are given in cm^-1^.

**Figure 7:**
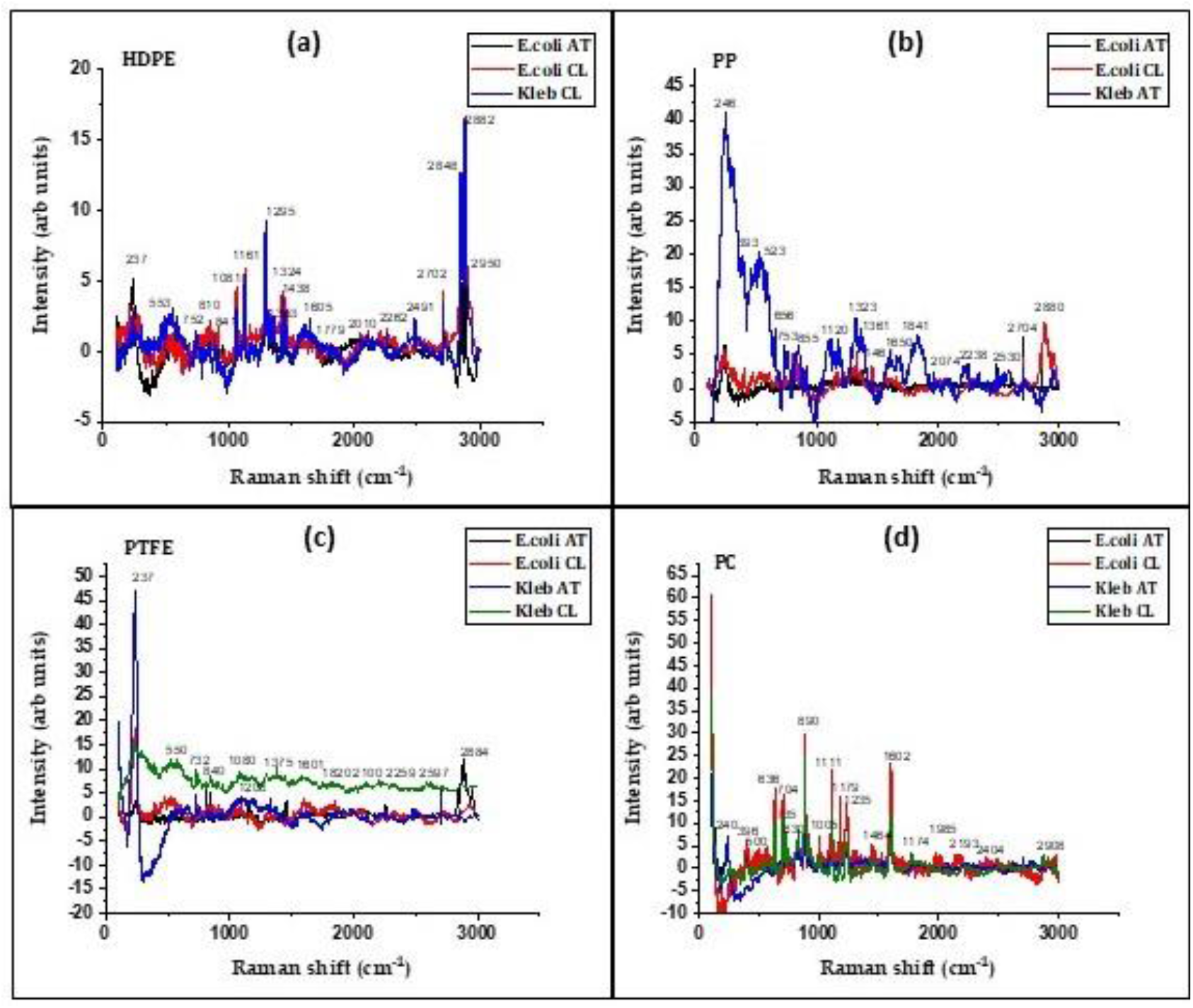
Surface Enhanced Raman Spectra of the bacterial biofilms. *Klebsiella pneumoniae* ATCC and Clinical, and *Escherichia coli* ATCC and Clinical bacterial biofilms grown on (a) HDPE, (b) PP, (c) PTFE, and (d) PC. Prominent peak positions are given in cm^-1^.

**Table 1.**
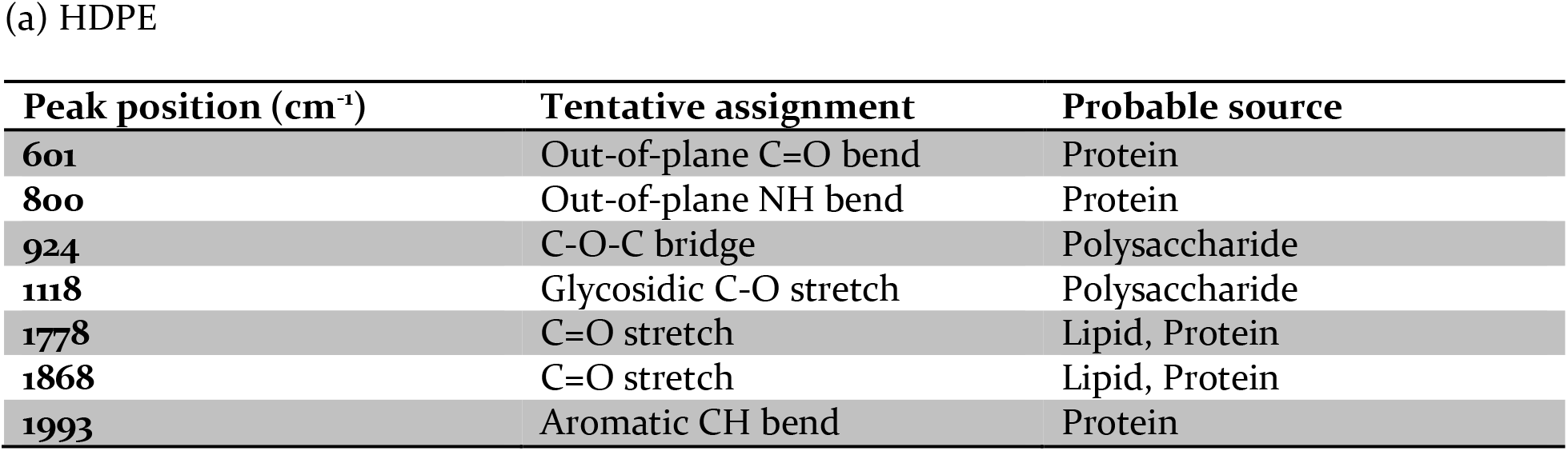
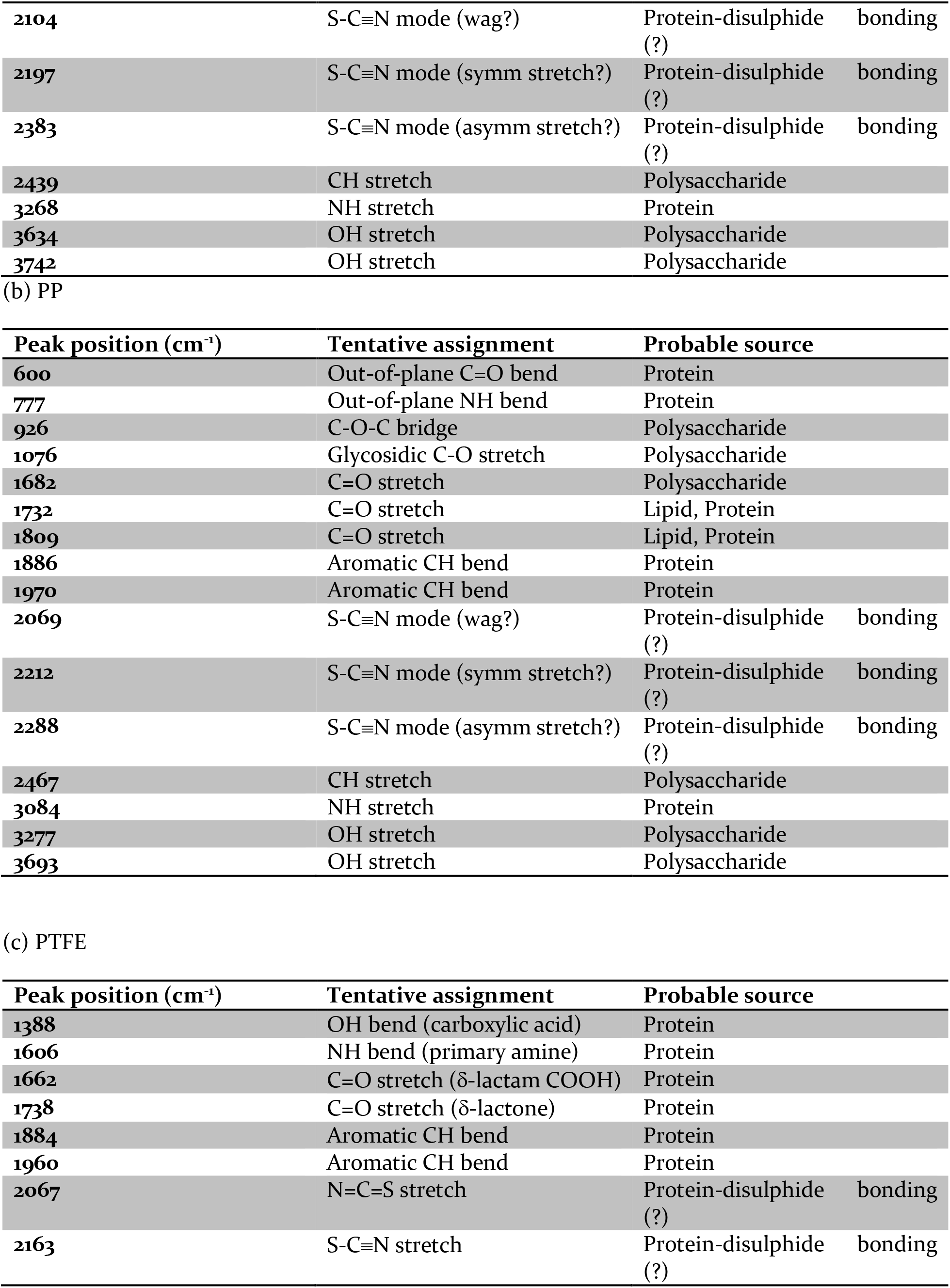
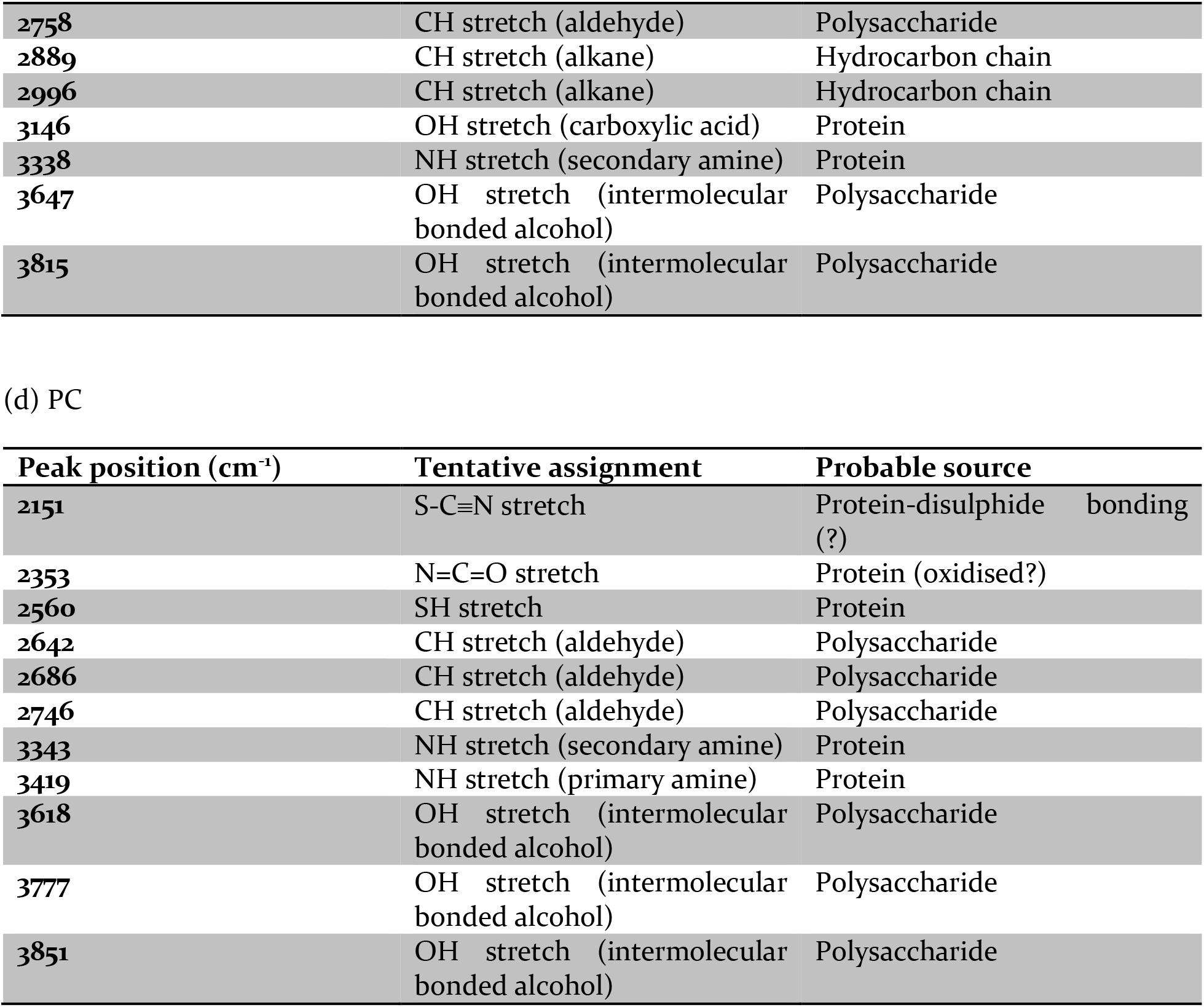
Fourier Transform Infrared Spectra of *K. pneumoniae* ATCC, *K. pneumoniae* Clinical, *E. coli* ATCC and *E. coli* Clinical bacterial biofilms grown on BSA-coated

**Table 2.** Surface Enhanced Raman Spectra of *K. pneumoniae* ATCC, *K. pneumoniae* Clinical, *E. coli* ATCC and *E. coli* Clinical bacterial biofilms grown on BSA-coated

| Peak position ($\text{cm}^{-1}$) | Tentative assignment | Probable source |
| --- | --- | --- |
| 553 | S-S stretch | Polysaccharide |
| 752 | Adenine ring deformation | Nucleic Acid |
| 810 | O-P-O stretch | Lipid |
| 841 | Tyrosine Fermi resonance | Protein |
| 1081 | C-O-C Glycosidic breathing | Polysaccharide |
| 1161 | Tyrosine C-N stretch | Protein |
| 1295 | C-O-H/H-C-C deformation | Polysaccharide |
| 1324 | C-H deformation | Protein |
| 1353 | C=C stretch | Lipid |
| **1438** | C-H<sub>2</sub>/CH<sub>3</sub> deformation | Lipid |
| **1605** | C-COOH stretch | Protein |

(b) PP
| **Peak position (cm<sup>-1</sup>)** | **Tentative assignment** | **Probable source** |
| --- | --- | --- |
| **393** |  | Polysaccharide |
| **523** | S-S stretch | Polysaccharide |
| **656** | Tyrosine C-S stretch | Protein |
| **753** | Adenine ring deformation | Nucleic Acid |
| **855** | Tyrosine Fermi resonance | Protein |
| **1120** | C-O-C Glycosidic breathing | Polysaccharide |
| **1323** | C-H deformation | Protein |
| **1361** | C=C stretch | Lipid |
| **1461** | C-H<sub>2</sub>/CH<sub>3</sub> deformation | Lipid |
| **1650** | Amide | Protein |

(c) PTFE
| **Peak position (cm<sup>-1</sup>)** | **Tentative assignment** | **Probable source** |
| --- | --- | --- |
| **550** | S-S stretch | Polysaccharide |
| **732** | Adenine | Nucleic Acid |
| **840** | Tyrosine Fermi resonance | Protein |
| **1080** | C-O-C Glycosidic breathing | Polysaccharide |
| **1203** | Phenylalanine | Protein |
| **1375** | C=C stretch | Lipid |
| **1601** | C-COOH | Protein |

(d) PC
| **Peak position (cm<sup>-1</sup>)** | **Tentative assignment** | **Probable source** |
| --- | --- | --- |
| **396** |  | Polysaccharide |
| **500** | S-S stretch | Polysaccharide |
| **636** | Tyrosine C-S stretch | Protein |
| **704** |  | Protein |
| **735** | Adenine ring deformation | Nucleic Acid |
| **830** | Tyrosine Fermi resonance | Protein |
| **890** |  | Protein |
| **1005** | Tryptophan ring breathing | Protein |
| **1111** | Tyrosine C-N stretch | Protein |
| **1179** | C-C stretch | Carotenoid |
| **1235** | Amide III | Protein |
| **1438** | C-H<sub>2</sub>/CH<sub>3</sub> deformation | Lipid |
| **1605** | C-COOH stretch | Protein |

### 3.3 Discussion

The above observations suggest that the EPS of the biofilm, secreted by the bacteria is generated to interact with the hydrocarbon and dipolar/dissociable moieties of the BSA-covered substrates to reduce the surface free energy for stable growth of the biofilm. As discussed in section 2.3, the distribution of these moieties on a BSA-covered polymer surface is decided by the specific polymer-BSA interaction. Thus, it is expected that the EPS of a biofilm on a particular BSA-covered polymer surface will also be largely decided by the polymer-BSA interaction of that surface. It is also interesting to note that the EPS on BSA-HDPE and on BSA-PP are almost identical and polysaccharide-rich, - consistent with the homologous composition of these polymers. The EPS on BSA-PTFE is protein-rich whereas that on BSA-PC has close proportions of polysaccharides and proteins.

**Figure 8.**
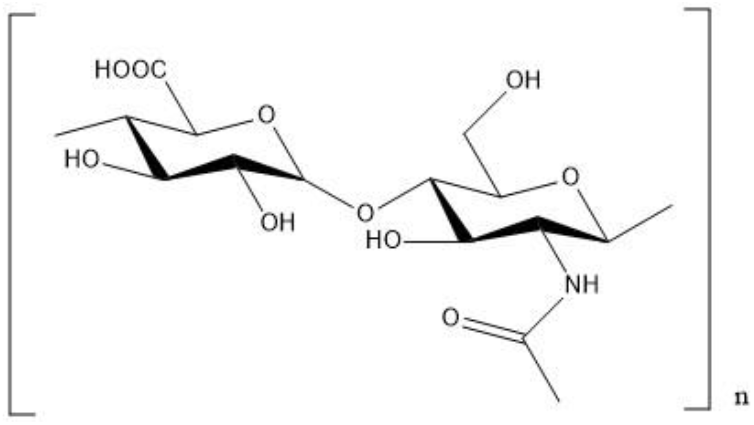
Molecular structure of a generic polysaccharide.

Inspection of the generic structure of polysaccharides (Figure 8) richer in hydroxyl groups than proteins (richer in the carboxyl and amine groups) may give some idea as to the specific interactions involved in producing the EPS. In absence of spatially resolved spectroscopic data a clearer picture is not available at this stage and work towards that end is underway.

## 4. Conclusion and Outlook

We have carried out Contact Angle Measurements, Scanning Electron Microscopy, Fourier Transform Infrared Spectroscopy, and Surface Enhanced Raman Spectroscopy of biofilms of *K. pneumoniae* and *E. coli* (ATCC and Clinical strains) on polymer substrates - HDPE, PTFE, PP, and PC which had been previously adsorbed with BSA for 24 hours. These studies prove conclusively that the BSA-polymer interaction decides the wetting properties, the intermolecular forces and the composition of the EPS of the bacterial biofilms, irrespective of the strain and type of bacteria involved.

These results are to be extended to gram positive bacteria and *in vivo* studies for understanding the specific nature of the BSA-polymer interaction and its role in the growth of the EPS. They form the first step towards the final goals of elimination of clinically pathogenic biofilms from physiological environments through noninvasive methods. Such studies will also have the potential for developing suitable biomaterial surfaces for negation of biofilm formation and may be extended for the enhancement of beneficial biofilms for widespread applications in environmental protection.

## Acknowledgements

The research has been carried out with the support of The Royal Society Grant (NIF\R1\181300). S DuttaSinha is thankful to The Royal Society, UK and Science and Engineering research Board (SERB), India for granting a Newton International Fellowship. AD would like to thank the Department of Atomic Energy, Government of India for a Raja Ramanna Fellowship, the Director, CSIR-Central Glass and Ceramic Research Institute for hosting the position of Emeritus Scientist, and Swapnasopan Datta of Jawaharlal Nehru Centre for Advanced Scientific Research for the figures of the molecular structures

## Data availability statement

The raw/processed data required to reproduce these findings cannot be shared at this time as the data also forms part of an ongoing study.

